# Persistent Elevation of Electrical Pain Threshold following Continuous Theta Burst Stimulation over Primary Somatosensory Cortex in Humans

**DOI:** 10.1101/724344

**Authors:** Nishant Rao, Yen-Ting Chen, Regan Ramirez, John Tran, Sheng Li, Pranav J. Parikh

## Abstract

**Background:** Primary somatosensory cortex (S1) is involved in pain processing and thus its suppression using neuromodulatory techniques such as continuous theta burst stimulation (cTBS) might be a potential pain management strategy in patients with neuropathic pain. S1 cTBS is known to elevate pain threshold in young adults. However, the persistence of this effect is unknown.

**Objective/Hypothesis:** We hypothesized persistent elevation of pain threshold following cTBS over S1 in healthy, young adults.

**Methods:** We recruited ten subjects in a sham-controlled crossover design and recorded their electrical pain threshold (EPT) for 40 min following cTBS over S1. We assessed corticospinal excitability (CSE) to rule out the involvement of primary motor cortex due to spread of current.

**Results:** cTBS over S1 elevated EPT without modulating CSE compared to sham stimulation. EPT was elevated for 40 min post-cTBS.

**Conclusions:** S1 can be focally targeted using cTBS for a longer lasting pain relief in patients.

## INTRODUCTION

Understanding cortical mechanisms for pain processing has been of considerable clinical interest for design of effective pain management strategies using neuromodulatory techniques [1–6]. Several neuroimaging studies have suggested involvement of cortical areas such as dorsolateral prefrontal cortex, primary somatosensory cortex, secondary somatosensory cortex, and primary motor cortex in processing of painful stimuli [1,7–10]. Transcranial magnetic stimulation (TMS) has been used to test the critical role of these areas in pain processing [2,3,8,11–13]. TMS-induced disruption of primary somatosensory cortex (S1) has been shown to increase reports of pain and elevate pain threshold in able-bodied individuals, thus suggesting the causal involvement of S1 in pain processing [2,14]. For instance, continuous theta burst stimulation (cTBS; a form of repetitive TMS), over S1 significantly reduced the perception of CO_2_ laser-evoked painful stimuli delivered to the contralateral hand when compared to the ipsilateral hand assessed immediately after S1 stimulation [2]. However, the persistence of after effects of S1 stimulation on pain threshold remains unknown.

Moreover, due to proximity of S1 and primary motor cortex (M1) regions, TMS over S1 could also induce current in the surrounding M1 region [15]. As M1 is also known to be critically involved in pain processing [4,5,7,11], the finding of elevated pain threshold reported following TMS over S1 might be confounded by changes in M1 excitability. Therefore, whether S1 plays a role in pain processing independent of M1 remains unknown. We studied the time course of the effects of cTBS delivered to S1 on electrical pain threshold (EPT), in healthy young adults, in a sham-controlled, crossover design. We hypothesized that cTBS over S1 would elevate EPT in healthy young adults. Because cTBS over M1 is known to reduce corticospinal excitability (CSE) for at least 30 min [16–18], we expected that the effects of cTBS over S1 on EPT will last for at least 30 minutes. We addressed the possibility of current spread to M1 by assessing CSE with single TMS pulses delivered over M1 [18,19].

## MATERIALS AND METHODS

Ten healthy, young, right-handed subjects [20] (mean±SD: 25.30±4.81 years; 4 females) provided written informed consent to participate in two sessions in a counterbalanced order. We estimated the effect size and sample size using our published data [21] on the change in electrical pain threshold (EPT) following breathing-controlled electrical stimulation in young adults [21–23]. To achieve at least 90% power to detect change in EPT following cTBS over S1, a sample size of 10 would be required assuming a Type I error of 0.05 and an effect size of 0.32 [24]. This sample size was consistent with that used in previous studies [17,21,25,26]. The study was approved by the Institutional Review Board of the University of Houston.

Electrical pain threshold (EPT) was measured over the abductor pollicis brevis (APB) muscle. Three repetitions were performed, and the corresponding average was used (see *SI* for details). We assessed CSE by measuring the size of motor evoked potentials (MEP) elicited in the first dorsal interosseous muscle (FDI) of the right hand across 10 trials. Tactile sensitivity and electrical sensory threshold were also assessed (see *SI*). We assessed each measure once before cTBS (PRE) and then every 10 min starting immediately post cTBS (POST_0_) until 40 min (POST_10_, POST_20_, POST_30_, POST_40_).

We used cTBS [16,17,27] to disrupt left S1 representing the contralateral hand. To target S1 (cTBS_S1_), we positioned the TMS coil over postcentral gyrus posterior to the M1 FDI hotspot identified on subject-specific MRI scans [17,28]. For sham stimulation (cTBS_SHAM_), same stimulation parameters were used, but the coil was placed perpendicular over the left S1 region so that no relevant current flow was induced in the cortical tissue [29–31].

We used repeated measures analysis of variance (α=0.05) with within-subject factors of CONDITION (cTBS_S1_, cTBS_SHAM_) and TIME (PRE, POST_0_, POST_10_, POST_20_, POST_30_, POST_40_). Posthoc paired t-tests were corrected for multiple comparisons using the false discovery rate at p<0.05 [32,33].

## RESULTS

No subjects reported any side effects during or after the experimental sessions.

EPT increased after cTBS_S1_ as compared to cTBS_SHAM_ (CONDITION×TIME interaction: F_5,45_=3.37, p=0.011, η_p_^2^=0.27; **Fig.1A**). Following cTBS_S1_, EPT increased from PRE to Post_0_ (t_9_=3.56, p=0.006), Post_10_ (t_9_=2.43, p=0.038), Post_20_ (t_9_=2.85, p=0.019), Post_30_ (t_9_=3.12, p=0.012), and Post_40_ (t_9_=2.96, p=0.016). There was no change in EPT following cTBS_SHAM_ (all t_9_<1.57, all p>0.15; refer **Figs. 1C** and **1D** for subject-wise changes in EPT). There was no difference in PRE EPT between cTBS_S1_ and cTBS_SHAM_ (t_9_=0.68, p=0.51). The results for tactile sensitivity and electrical sensory threshold are reported in *SI*.

**Figure 1.**
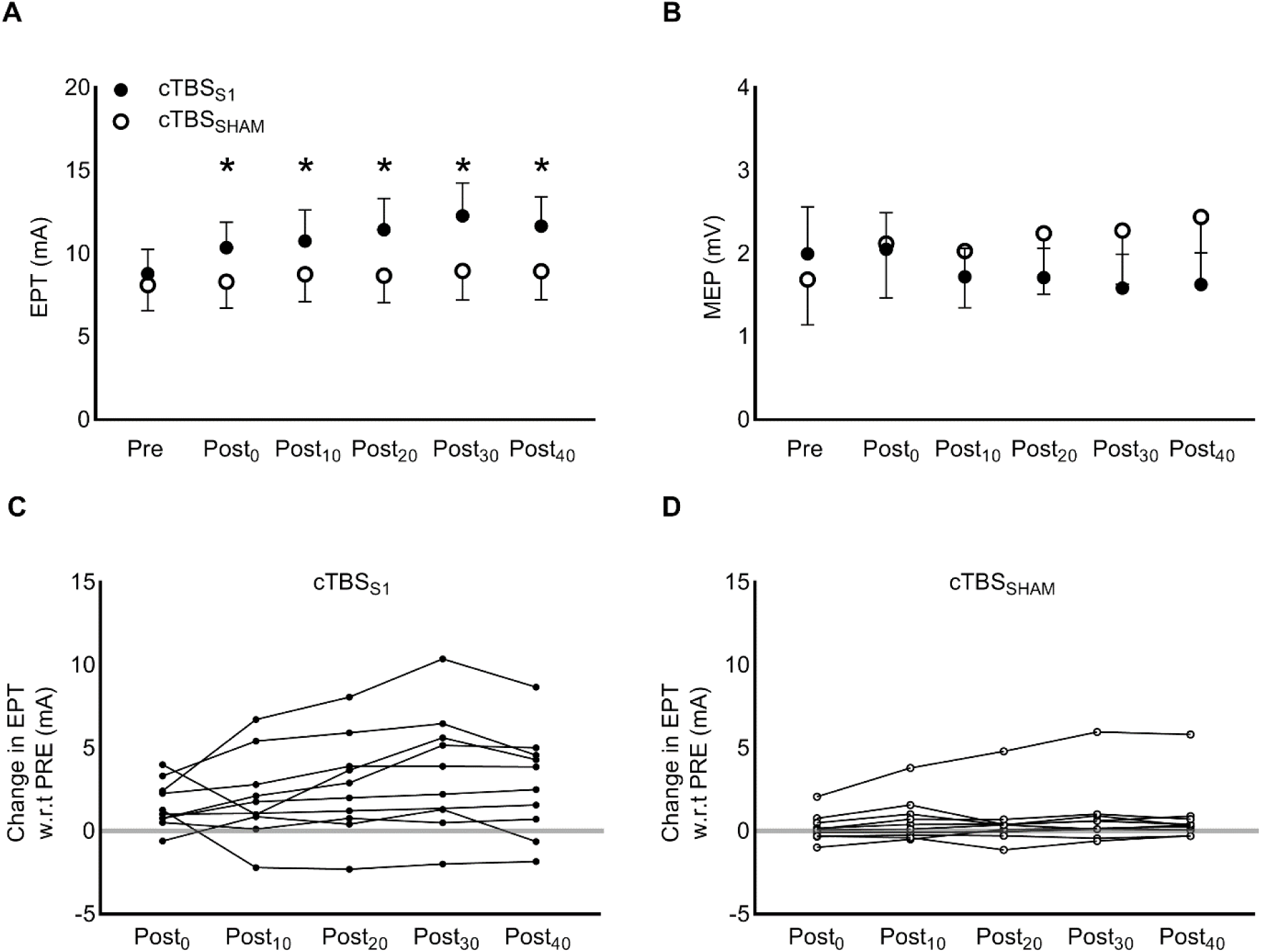
Time course of electrical pain threshold (EPT) and corticospinal excitability (CSE) post cTBS over S1. **A.** Increase in EPT following cTBS over S1 compared to sham stimulation (significant *Condition* × *Time* interaction: p=0.011). Asterisks indicate a significant increase in EPT at Post time point with respect to PRE (p < 0.05, FDR-corrected). **B.** No change in group-level data for MEP, a measure of CSE, assessed over M1 following cTBS_S1_ and cTBS_SHAM_. For **A** and **B**, each circle and error bar represent mean and standard error across subjects (n=10) respectively. **C.** Subject-wise time course of change in EPT with respect to PRE following cTBS over S1 (solid circles). **D.** Subject-wise time course of change in EPT with respect to PRE following sham stimulation (open circles). In addition to EPT and CSE, we also measured tactile sensitivity and electrical sensory thresholds as our secondary measures and these findings are presented in *Supplementary Information*.

We found no difference in MEP amplitude across cTBS_S1_ and cTBS_SHAM_ (no CONDITION×TIME interaction: F_5,45_=2.24, p=0.07, η_p_^2^=0.19; no CONDITION effect: F_1,9_=1.39, p=0.27 η_p_^2^=0.13; no TIME effect: F_5.45,_=0.23, p=0.95, η_p_^2^=0.03; **Fig.1B**). There was no difference in PRE MEP amplitude between cTBS_S1_ and cTBS_SHAM_ (t_9_=1.6, p=0.14). These findings indicate that the stimulation current delivered over S1 did not affect CSE.

## DISCUSSION

We found that cTBS over S1 elevated EPT for 40 min in healthy young adults without changes in the MEP size assessed over M1. A spread of current from S1 to M1 would have influenced MEP size because previous work including ours have found a reduction in MEP size following disruption of M1 activity using cTBS at rest [16–18]. This finding of unchanged MEP size precludes the role of M1 in the observed effects of cTBS over S1 on EPT. Our findings provide evidence that a single session of cTBS over S1 could provide persistent analgesic effect in healthy adults. In contrast, previous studies only reported analgesic effects immediately [2] or 5 minutes following cTBS over S1 [8]. In summary, our findings suggest that cTBS over S1 can serve as a potential focal neuromodulatory tool for pain management. The long-term effect after repetitive use of this intervention and its application in clinical populations need further investigation.

## ACKNOWLEDGMENT

This study was partially supported by the Core for Advanced MRI (CAMRI) pilot grant from Baylor College of Medicine and the High Priority Area Research Seed Grant from the University of Houston to PP.

## SUPPLEMENTARY INFORMATION

### METHODS

#### Assessment of corticospinal excitability (CSE)

We measured CSE by assessing the size of motor evoked potentials (MEP) elicited in the first dorsal interosseous muscle (FDI) of the right hand. The FDI muscle activity was recorded using differential surface electrodes (Delsys Bagnoli EMG System, Boston, MA). The data were sampled at 5 kHz using CED data acquisition board (Micro1401, Cambridge, England).

Single-pulse TMS was used to assess CSE over primary motor cortex (M1) during the experiment [1,2]. We first estimated the resting motor threshold (rMT) by delivering suprathreshold single monophasic TMS pulses (Magstim 200, Whitland, UK). The TMS coil was held tangential to the scalp and perpendicular to the presumed direction of the central sulcus, 45° from the midsagittal line, with the handle pointing backward, inducing current in the posteroanterior direction. The coil position was adjusted to optimize the motor-evoked potential (MEP) in the FDI muscle. Following this procedure, the rMT was estimated as the minimum TMS-intensity to elicit motor evoked potential (MEP) with an amplitude of ∼50 µV (peak-to-peak) for at least 5 of the 10 consecutive trials in the FDI muscle [1–4]. The TMS coil was stabilized using a coil holder mounted on the TMS chair (Rogue Research). The TMS coil was traced on the subject’s scalp using a surgical marker pen. The coil location was regularly checked for any displacement that might have occurred during a session. The average rMT across subjects (mean±SE) was 54±3% of the maximum stimulator output. The corticospinal excitability was assessed with the intensity set at 120% of rMT over the identified FDI region and averaged across 10 consecutive trials.

In addition, we also assessed active motor threshold (aMT) to set the stimulation intensity of continuous theta burst stimulation. For aMT estimation, subjects were instructed to exert 20% of individual’s maximum voluntary force (MVF) on a grip device instrumented with force transducers (Nano-25; ATI Industrial Automation, Garner, NC, 1 kHz sampling rate) with the tips of index finger and thumb using visual feedback provided on a computer monitor. Each subject was instructed to grip the device as hard as possible for one second followed by a break (∼1 min). This procedure was repeated three times, and we used the largest grip force as the MVF recorded across three trials. The aMT was determined as the TMS intensity that induced 200 μV peak-to-peak MEPs in 5 of 10 trials in the FDI muscle during force production at 20% of MVF [5]. The aMT was 44±2% (mean across all subjects±SE; n=10) of the maximum stimulator output.

#### Continuous theta burst stimulation

We used continuous theta burst stimulation (cTBS) to disrupt left primary somatosensory region representing the contralateral hand. Prior to the cTBS procedure, we obtained a high-resolution T1-weighted MRI scan (3T Philips Ingenia scanner) for each subject. A three-dimensional brain was reconstructed from the MRI slices to display the cortical surface (Brainsight software, Rogue Research Inc., Canada). For S1 cTBS, we positioned the TMS coil over the postcentral gyrus posterior to the M1 FDI hotspot [6]. The mean Montreal Neurological Institute coordinates of the stimulation sites for left S1 were −41.67±8.90, −28.27±6.57, 65.10±11.06 (x, y, z, mean ±SD; n=10).

We delivered cTBS over the left S1 using a figure-of-eight coil at 80% of aMT to temporarily disrupt its activity. Repetitive biphasic TMS pulses were delivered in the form of bursts of three pulses at 50 Hz at a rate of 5 Hz, i.e. 200 ms inter-burst interval, for 40 s. The cTBS protocol resulted in the delivery of 600 pulses [7]. The exact positioning of the coil was visually monitored throughout the stimulation duration. For the sham stimulation (cTBS_SHAM_), the same stimulation parameters were used, but the coil was placed perpendicular over the left S1 region so that no relevant current flow was induced in the cortical tissue [8,9]. The intensity of stimulation was well within the safety guidelines for TMS use, and consistent with that used by other TMS groups [7,10–15].

#### Tactile Sensitivity and Electrical Sensory Threshold Assessments

Tactile sensitivity (TS) was measured using Semmes-Weinstein Monofilaments Examination (SWME, Smith and Nephew Roland, Menominee Falls, WI) [16–19]. Tactile sensibility thresholds were obtained from the distal volar pads of the index finger. The index finger was tested approximately midway between the center of the pad and the radial margin of the finger. A threshold was recorded for the smallest filament diameter (buckling force in mg, according to the manufacturer’s calibration) that could be perceived on at least 70% of its applications.

Electrical sensory threshold (EST) and electrical pain threshold (EPT) were measured using manual triggering of electrical stimulator (DS7A; Digitimer, Hertfordshire, UK) via a surface bar electrode placed over the abductor pollicis brevis (APB) muscle. For EST, the intensity of electrical stimulation was started from zero and gradually increased in steps of 0.1 mA until the subject explicitly felt electrical stimulation. This was followed by EPT measurement. The intensity was started from the EST and increased in steps of 1 mA until the subject first felt the electrical stimulation to be painful. Participants were explicitly instructed that the aim of the study was not to assess maximum pain they can bear but to measure only their pain threshold. To improve consistency among subjects, they were advised to report a stimulus to be painful upon experiencing the pain level equivalent to 1 on the 0–10 visual analog scale [18]. Three repetitions were made for each measure and the corresponding average was used for both EST and EPT.

### RESULTS

TS reduced following cTBS_S1_ (main effect of TIME for nonparametric Friedman test, χ^2^ _5,10_=15.174, p=0.010; **Fig. S1A**). Posthoc comparisons were conducted using Wilcoxon signed-rank test and the change in TS from PRE to Post_0_ (Z=−1.826, p=0.068), Post_10_ (Z=−2.21, p=0.02), Post_20_ (Z=−2.21, p=0.02), POST_30_ (Z=−2.032, p=0.042) and POST_40_ (Z=−1.841, p=0.066) failed to reach the FDR corrected significance level. We did not find a change in TS following cTBS_SHAM_ (no main effect of TIME, χ^2^_5,10_ =8.582, p=0.127). There was no difference in PRE TS measure between cTBS_S1_ and cTBS_SHAM_ (Wilcoxon signed-rank test: Z=−1.000, p=0.317). The reduction in TS following cTBS_S1_ but not cTBS_SHAM_ is consistent with previous study [20–22] indicating the role of S1 in processing sensory information.

PRE EST measure was not different between cTBS_S1_ and cTBS_SHAM_ (t_9_=0.203, p=0.844). We observed a significant increase in EST following stimulation (main effect of TIME: F_5,45_=7.112, p<0.001, η_p_^2^=0.441). However, this increase was similar following cTBS_S1_ and cTBS_SHAM_ (no CONDITION×TIME interaction: F_5,45_=1.378, p=0.250, η_p_^2^=0.133; no main effect of CONDITION: F_1,9_=0.493, p=0.500, η_p_^2^=0.052; **Fig. S1B**). The increase in EST following cTBS_SHAM_ suggests habituation to repeated low intensity electrical stimulation of the thenar eminence [23]. It is likely that similar habituation to low intensity electrical stimulation was present in the cTBS_S1_ session and that it might have confounded the effects of cTBS over S1 on EST. Therefore, the EST measure, at least in our study, was not a reliable measure to study the effects of cTBS over S1 on sensory perception. It is important to note that similar habituation was not observed for other experimental measures such as EPT and TS.

---

**Figure S1.**
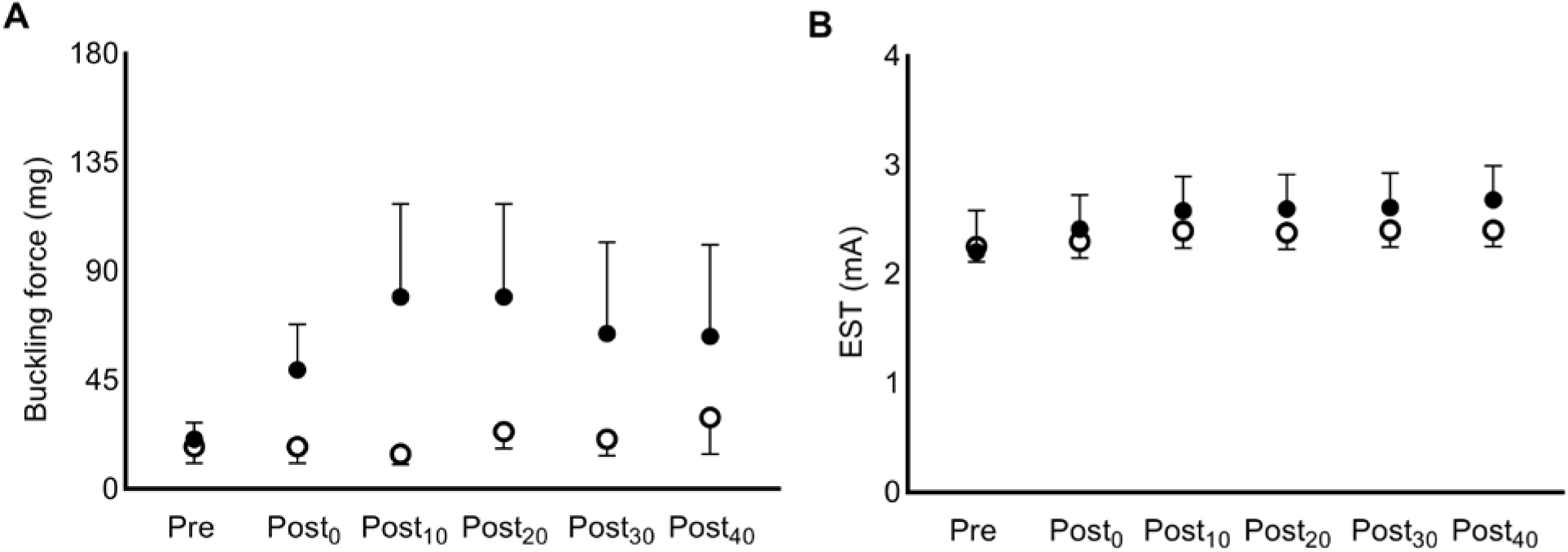
Time course of tactile sensitivity and electrical sensory threshold following cTBS. **A.** Tactile sensitivity threshold was recorded for the smallest filament diameter (buckling force in mg) that could be perceived on at least 70% of its applications. An increase in buckling force post cTBS_S1_ corresponds to a reduction in TS. There was no change in tactile sensitivity following sham stimulation (cTBS_SHAM_). **B.** Temporal drift in electrical sensory threshold following both cTBS_S1_ and cTBS_SHAM_. Each circle and error bar represent mean and standard error across subjects (n=10).

